# Morphine and methamphetamine trigger divergent post-transcriptional neuroimmune landscapes in the dorsal striatum

**DOI:** 10.64898/2026.04.01.716002

**Authors:** Alexander V. Margetts, Lauren L. Bystrom, Samara J. Vilca, Luis M. Tuesta

## Abstract

Opioid and methamphetamine use disorders (OUD and MUD) are characterized by enduring neural adaptations within brain reward circuitry, yet the cell-type-specific post-transcriptional mechanisms underlying these changes remain poorly understood. While microglia are essential for maintaining central nervous system homeostasis and modulating neuroinflammatory responses to drugs of abuse, their alternative splicing (AS) programs have not been defined in the context of addiction. This study characterized the microglial AS landscape in the mouse dorsal striatum during morphine and methamphetamine intravenous self-administration (IVSA), as well as following a 21-day period of abstinence. Analysis of RNA-sequencing data using rMATS and DEXSeq revealed that both drugs significantly dysregulate core splicing machinery, with skipped exons (SE) emerging as the most prevalent splicing event. Notably, morphine exposure induced a robust persistent splicing signature, comprising 736 exonic regions in 221 genes that remained altered through abstinence, whereas methamphetamine-induced changes were primarily reversible. Functional annotation predicted that approximately 27.5% of these events induce frameshifts, potentially impacting critical microglial pathways such as autophagy (*Wdr81*), chromatin remodeling (*Chd4, Kmt2c*), and RNA processing (*Hnrnpl, Mbnl2, Tia1*). These findings identify previously unrecognized post-transcriptional neuroimmune mechanisms and suggest that persistent splicing dysregulation in microglia may contribute to the long-term pathophysiology of OUD.

**Graphical Abstract:** 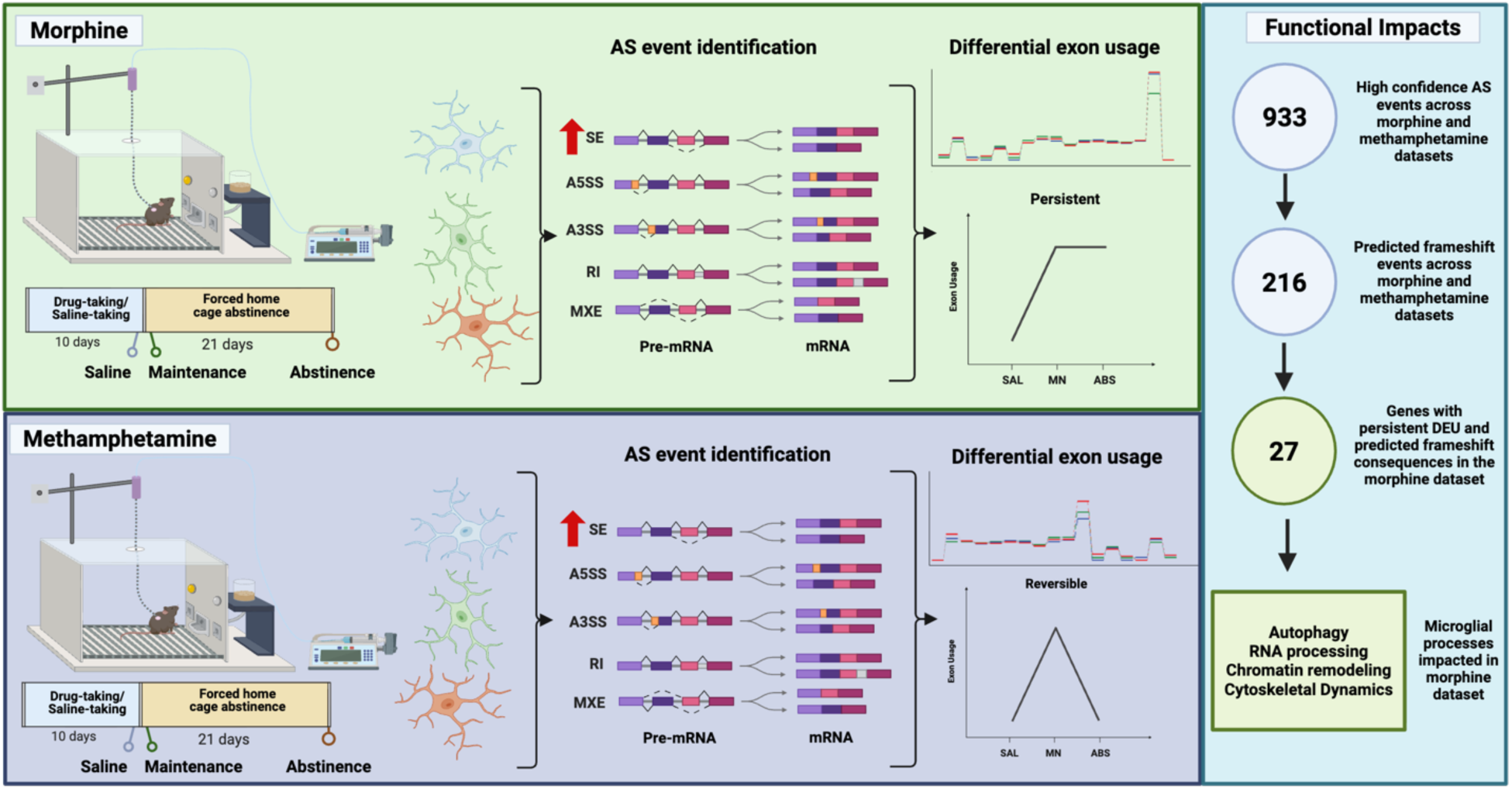

## Introduction

Opioid use disorder (OUD) remains a major public health crisis in the United States, producing profound mortality, morbidity, and economic costs. In 2017, an estimated 2.1 million Americans met criteria for past-year OUD, opioid overdose deaths exceeded 47,000, and the overall economic burden of OUD and fatal opioid overdose was estimated at approximately $1.02 trillion^1^. These trends underscore the need to define molecular mechanisms within brain reward circuitry that contribute to compulsive drug taking^2–4^. Methamphetamine use disorder (MUD) has likewise re-emerged as a public health problem in the United States. National estimates indicate that past-year methamphetamine use increased from approximately 1.4 million individuals in 2016 to 2.0 million in 2019, with roughly 1 million people meeting criteria for MUD^5^. MUD is associated with substantial psychiatric, cardiovascular, cerebrovascular, infectious, and neurocognitive comorbidities, yet there remain no FDA-approved pharmacotherapies for MUD^6,7^. Together, these observations establish MUD and OUD as critical areas for research.

Microglia are the resident immune cells of the CNS. Under homeostatic conditions, microglia continually surveil the neural environment, phagocytosing debris (e.g., dead cells, myelin, bacteria), participate in synaptic pruning, circuit refinement, and contribute to homeostatic support for neurons, glia, and vascular cell function^8,9^. Under disease or injury conditions, these same cells can adopt inflammatory phenotypes that shape neuroimmune signaling, synaptic remodeling, and neuronal survival^10^. Importantly, many classes of drugs can affect the neural microenvironment, especially in terms of neuroinflammatory processes^11–16^. It is believed that opioids can activate microglial inflammatory pathways through mechanisms involving TLR4/MD2 and downstream production of pro-inflammatory mediators such as TNF-α, IL-1β, and IL-6^17–20^. Methamphetamine similarly alters microglial function through innate immune and purinergic pathways, including TLR4- and P2X7R-dependent signaling, and may produce neuroinflammatory changes in reward-related regions^17,21–23^. These findings support the concept that microglia are active participants in the response to both opioid and methamphetamine exposure. Specifically, microglia of the dorsal striatum are critical to the overall function of the region which is involved in stimulus learning and habit formation as well as contributing to motor control^24^. Both opioids and psychostimulants are known to impact signaling in the dorsal striatum (hereby referred to as “striatum”)^25,26^, and as such, striatal circuitry has been a central research focus for the pathophysiology of substance use disorders (SUDs)^27,28^. Beyond neuronal adaptations, microglia can influence synaptic strength, inflammatory processes, and dopaminergic function, thereby shaping the circuit consequences of repeated drug exposure^27,29–33^.

Among the molecular mechanisms inducing long-lasting cellular adaptation, alternative splicing (AS) has emerged as an intriguing candidate. Recent evidence demonstrates that SUDs may alter AS programs across reward-related regions and identification of splicing changes separate from differential gene expression could point to post-transcriptional regulation. Importantly, cell-type-specific studies from other neurological disorders indicate that microglial AS programs are implicated in disease response. Specifically, in Alzheimer’s and multiple sclerosis disease models, microglia exhibit distinct AS signatures^34,35^, suggesting that microglial transcriptomes may be shaped by disease-associated splicing regulation. However, despite the interest in studies on AS dysregulation in neurodegenerative diseases, microglia specific AS events have not been examined in the context of SUDs.

To date, no study has defined cell-type-specific AS programs in microglia under opioid or methamphetamine self-administration conditions. Using morphine intravenous self-administration (IVSA) and publicly available data from methamphetamine IVSA, the present work sought to characterize the AS landscape of male murine striatal microglia following opioid and stimulant exposure. By identifying shared and distinct differential exon usage across these two addiction-relevant paradigms, this study has identified previously unrecognized post-transcriptional neuroimmune mechanisms that may contribute to OUD and MUD.

## Results

### Alternative splicing patterns in a morphine-taking and morphine-abstinent mice

The role of alternative splicing (AS) in microglia during OUD is not well characterized. Thus, we first set out to understand the transcriptional consequences of morphine-taking and morphine abstinence on splicing related machinery in striatal microglia. To investigate the role of AS in microglial responses to morphine-taking and abstinence, we utilized a model of morphine intravenous self-administration (IVSA) in mice. Briefly, mice were trained to earn intravenous infusions of morphine (0.1 mg/kg/infusion; 36 μl/infusion; 3 second infusion) or saline in an operant chamber for 15 consecutive days (morphine-taking (MN) and saline-taking (SAL)), and were then subjected to 21 consecutive days of forced home cage-abstinence (morphine abstinence (ABS)). Following self-administration (SAL and MN) and forced home cage abstinence (ABS), microglia were isolated from the striatum and RNA-sequencing was conducted to determine the transcriptional responses of drug-taking and forced abstinence. Gene ontology (GO) analysis revealed significant (padj < 0.05, qvalue < 0.1) enrichment of 741 differentially expressed genes (DEGs) (padj < 0.05) in 97 pathways related to AS mechanisms including spliceosome complex, RNA splicing, mRNA processing and regulation of mRNA processing across multiple pairwise comparisons (**Fig 1A**). DEGs in these pathways included core splicing components such as small nuclear ribonucleoprotein polypeptide (*Snrnp*) A1 (*Snrnpa1)*, B (*Snrnpb*), B2 (*Snrnpb2*) and 70 (*Snrnp70*) (**Supplementary Data File 1**)^36^. Moreover, genes involved in specific spliceosome components were significantly dysregulated between conditions. This includes the Sf3b complex, in which mutations have been associated with several human cancers^37^, and more specifically splicing factor 3B (*Sf3b*) subunit 1 (*Sf3b1*), subunit 2 (*Sf3b2*), subunit 3 (*Sf3b3*), subunit 4 (*Sf3b4*), and subunit 5 (*Sf3b5*) (**Supplementary Data File 1**).

**Figure 1.**
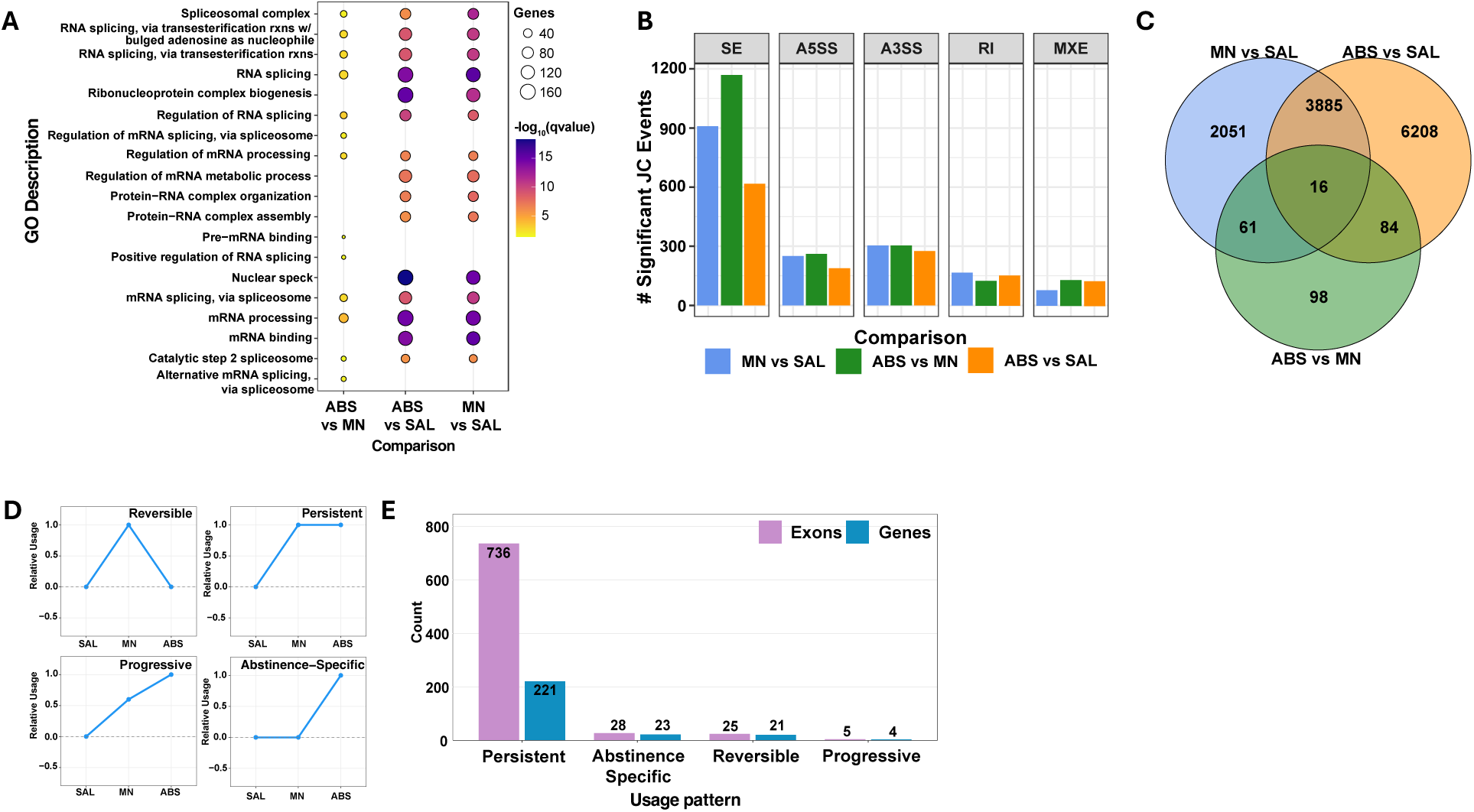
AS contributes to persistent changes in striatal microglia throughout morphine-taking and abstinence. (**A**) Selected GO terms (padj <0.05, qvalue < 0.1) related to splicing pathways in SAL, MN and ABS from striatal microglial differential gene expression after morphine IVSA. (**B**)Significant rMATS junction spanning (JC) counts of splicing events in striatal microglia throughout morphine IVSA (FDR < 0.05, |ΔPSI| > 0). (**C**) Venn-Diagram of significant differentially used exons across phases of morphine administration (FDR < 0.05, |L2FC| > 0.5). (**D**) Schematic classification of patterns for exon usage with example exon usage data. (**E**) Bar plot counts of high confidence exons (those with significant rMATS events in the parent gene (FDR < 0.05, |ΔPSI| > 0.5) and DEXSeq analyses (FDR < 0.05, |L2FC| > 0.5) and genes containing high confidence exons across identified usage patterns in morphine IVSA.

While differential gene expression analysis identified several dysregulated splicing components, we sought to characterize the mechanisms by which AS is impacted during morphine-taking and morphine abstinence. To this end, we employed rMATS^38^ to generate junction-spanning counts of AS events classified as skipped exon (SE), mutually exclusive exon (MXE), alternative 3’ splice site (A3SS), alternative 5’ splice site (A5SS), and retained intron (RI) (**Fig 1B**). Interestingly, we found that SE events were the most predominant across all pairwise comparisons, with almost 1,200 SE events occurring when comparing ABS vs MN, 900 SE events when comparing MN vs SAL and 600 when comparing ABS vs SAL (FDR < 0.05, |ΔPSI| > 0) (**Fig 1B**). A5SS, A3SS, RI and MXE events while present, contributed far less to the overall number of AS events (**Fig 1B**). To further understand how splicing contributes to changes in exon usage, we conducted differential exon usage (DEU) analysis using DEXSeq^39^ across SAL, MN and ABS phases of morphine IVSA in a pairwise manner. Of note, each pairwise comparison resulted in distinct exonic usage (2,051 exonic regions in MN vs SAL, 6,208 exonic regions in ABS vs SAL and 98 exonic regions in ABS vs MN) (FDR < 0.05 and |L2FC| > 0.5), and we identified a number of overlaps between comparisons, including 16 exonic regions that shared differential usage in all comparisons (FDR < 0.05 and |L2FC| > 0.5) (**Fig 1C**). To better characterize how these changes persist throughout multiple phases of morphine-taking and abstinence we selected for high-confidence AS changes (genes with rMATS-detected AS events and DEU based on FDR and L2FC significance) as reversible, persistent, progressive or abstinence-specific (**Fig 1D**). Persistent exon usage changes, those that occur during morphine-taking and continue to persist into abstinence, represented the largest classification with 736 exonic regions in 221 genes meeting those criteria (**Fig 1E**). While detection of other patterns such as abstinence-specific, reversible and progressive exon usage changes were present, they represented a smaller fraction (28 exonic regions in 23 genes, 25 exonic regions in 21 genes and 5 exonic regions in 4 genes, respectively) (**Fig 1E**).

Next, we mapped the 736 exons exhibiting persistent dysregulation to their respective genes and conducted GO analysis to determine which biological pathways may be altered by changes in exon splicing because of morphine-taking and abstinence. GO analysis revealed that genes exhibiting persistent exon usage dysregulation were enriched for pathways involved in epigenetic regulation (histone modification, chromatin organization, histone methylation, and histone lysine methylation) (**Fig 2A**), AS pathways (mRNA processing, RNA splicing, regulation of mRNA splicing, via spliceosome) (**Fig 2A**), and critical microglial cell functions including cytoskeletal restructuring and signaling (regulation of axon extension, regulation of developmental growth) (**Supplementary Data File 1**). Of interest, we found several critical RNA splicing genes to be altered by AS mechanisms, which may contribute to changes in splicing activity. More specifically, heterologous nuclear ribonucleoprotein L (*Hnrnpl*) contained a RI event and 4 persistent DEU’s (E3, E15, E16 and E19) (**Fig 2B**), muscleblind-like splicing regulator 2 (*Mbnl2*), contained 1 SE, 1 MXE, and 1 A3SS event along with 1 persistent DEU (E15) (**Fig 2C**), and T-cell intracellular antigen-1 (*Tia1*), contained 2 SE, 1 A3SS and 1 RI events as well as 1 persistent DEU (E25) (**Fig 2D**) (**Supplementary Data File 1**). *Hnrnpl*, *Mbnl2, and Tia1* are integral mediators of AS events^40–42^. Further, WD repeat domain 81 (*Wdr81*), which is a known regulator of protein aggregation and autophagy^43^, contained an A5SS event along with 2 persistent DEU’s (E1 and E2) (**Fig 2E**) (**Supplementary Data File 1**). Interestingly, OUD is known to induce changes in epigenetic regulation of gene expression^44,45^. Indeed, we identified key epigenetic regulator genes such as chromatin helicase DNA binding protein 4 (*Chd4*), which contained a RI event and 2 persistent DEUs (E4 and E42) (**Fig 2F**) and lysine methyltransferase 2c (*Kmt2c*), which contained 2 SE events, and 37 persistent DEUs (**Fig 2G**) (**Supplementary Data File 1**). These data indicate that morphine-taking induces pervasive AS changes in striatal microglia which may impact epigenetic regulation of gene expression as well as cellular functions critical to the homeostatic functions of microglia, which can persist for 21 days after the last exposure to the drug.

**Figure 2.**
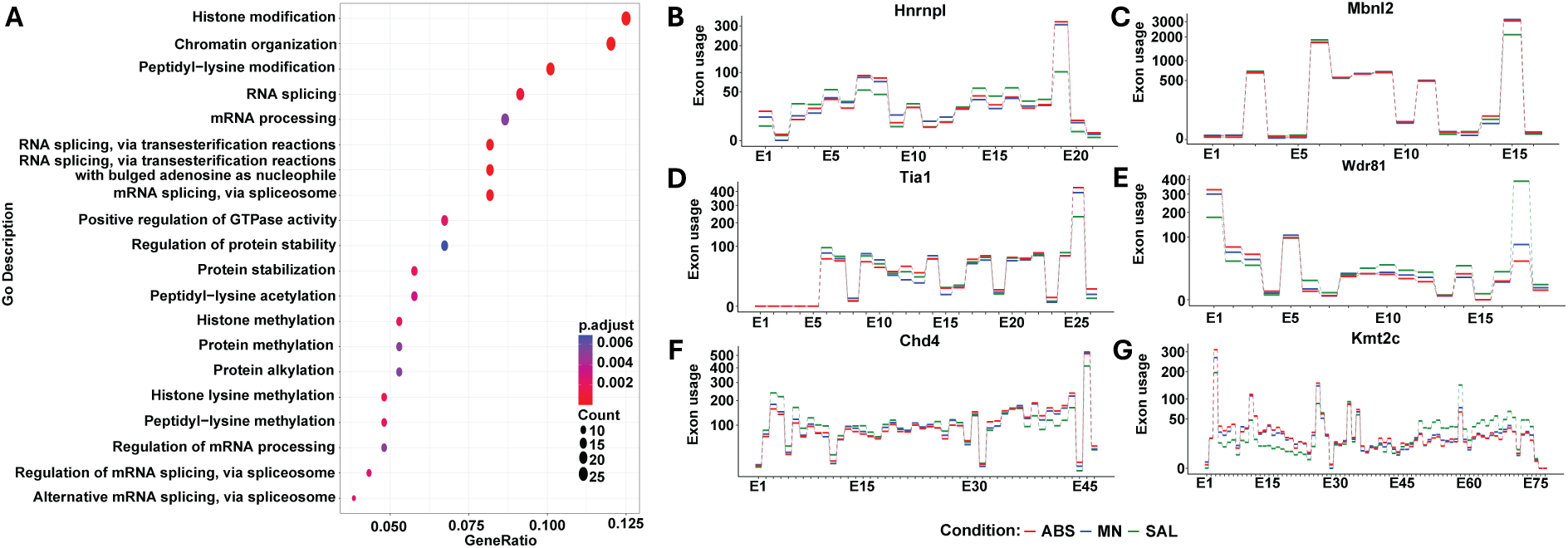
Pathway analysis of persistent AS events in morphine IVSA. **(A)** GO pathway analysis (padj <0.05, qvalue < 0.05) of genes containing an AS event and DEU’s classified as persisting throughout morphine abstinence. (**B-G**) Selected exon usage plots for exonic regions in genes from enriched GO pathways.

### Alternative splicing patterns in methamphetamine-taking and methamphetamine-abstinent mice

After identifying significant alterations in striatal microglia AS during morphine-taking and abstinence, we next sought to understand if these functional consequences were present in other SUDs. We therefore first examined the transcriptional consequences of methamphetamine-taking and abstinence on splicing related machinery in our model. To investigate the role of AS in microglia responses to methamphetamine-taking and abstinence, we utilized publicly available RNA-sequencing data (GSE245951) from a study of transcriptional consequences of methamphetamine IVSA on striatal microglia in male mice^46^. IVSA methods were carried out as described in *Vilca et al.*^46^ Briefly, mice were trained to earn intravenous infusions of methamphetamine (0.05 mg/kg/infusion; 14 μl/infusion; 2 second infusion) or saline in an operant chamber for 15 consecutive days (methamphetamine-taking (MN) and saline-taking (SAL)), and were then subjected to 21 consecutive days of forced home cage-abstinence (methamphetamine abstinence (ABS))^46^.Transcriptional data were regenerated to determine the transcriptional responses of drug-taking and forced abstinence. GO analysis on DEGs (padj < 0.05) revealed significant (padj < 0.05, qvalue < 0.1) enrichment of 49 DEGs in 23 pathways related to AS mechanisms including spliceosome complex, positive regulation of RNA splicing, negative regulation of RNA splicing and mRNA processing, notably only in the ABS vs MN and MN vs SAL groups (**Fig 3A**). The ABS vs SAL DEGs resulted in no significant enrichment of AS pathways, as defined herein. Similar to the findings from morphine-taking and abstinence, differential gene expression analysis conducted by *Vilca et al*. revealed that these pathways contain DEGs critical to splicing components including serine/arginine-rich splicing factor 3 (*Srsf3*), *Srsf5*, *Srsf7*, *Srsf7*, and *Srsf10* (**Supplementary Data File 1**). Moreover, genes involved in specific spliceosome components such as the SF3A complex, including splicing factor 3a subunit 3 (*Sf3a3*) was found to be significantly dysregulated between conditions, as noted by the study.

**Figure 3.**
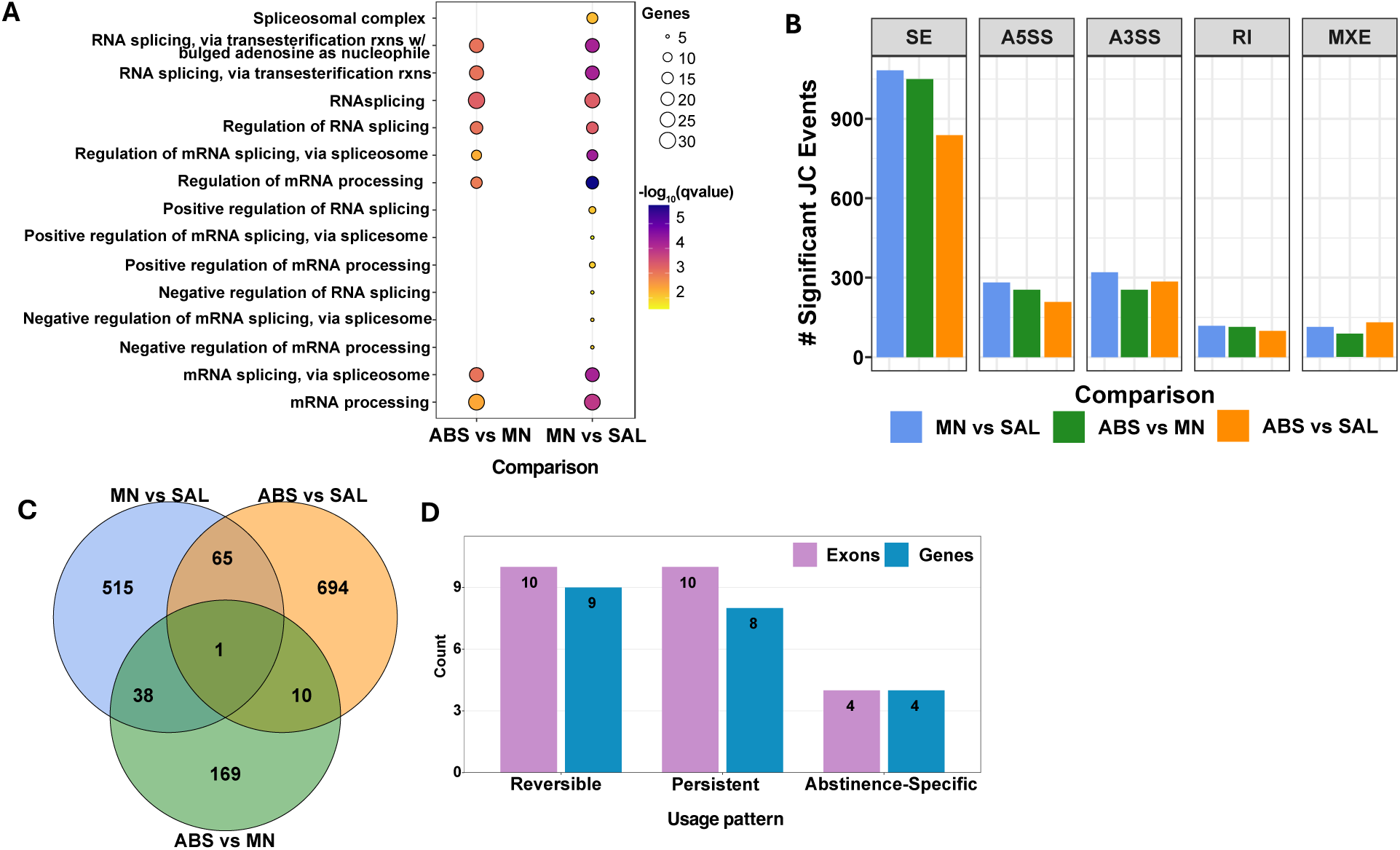
**AS contributes to persistent changes in striatal microglia throughout methamphetamine IVSA**. (**A**) Selected GO terms (padj <0.05, qvalue < 0.1) related to splicing pathways in SAL, MN and ABS from striatal microglial differential gene expression after methamphetamine IVSA. (**B**) rMATS junction spanning (JC) counts of splicing events in striatal microglia throughout methamphetamine IVSA (FDR < 0.05, |ΔPSI| > 0). (**C**) Venn-Diagram of significant differentially used exons across phases of methamphetamine administration (FDR < 0.05, |L2FC| > 0.5). (**E**) Bar plot counts of high confidence exons (those with significant rMATS events in the parent gene (FDR < 0.05, |ΔPSI| > 0.5) and DEXSeq analyses (FDR < 0.05, |L2FC| > 0.5) and genes high containing confidence exons across identified expression patterns in methamphetamine IVSA.

While differential gene expression analysis identified several dysregulated splicing components, we sought to characterize the mechanisms by which AS is impacted during methamphetamine-taking and abstinence. To do this, we again employed rMATS^38^ to generate junction-spanning counts of AS events as classified previously (**Fig 3B**). Interestingly, we again found that SE events were the most predominant across all pairwise comparisons, with greater than 1,000 SE events occurring when comparing ABS vs MN and MN vs SAL and more than 750 when comparing ABS vs SAL (FDR < 0.05, |ΔPSI| > 0) (**Fig 3B**). Compared to morphine, methamphetamine-taking as well as abstinence relative to saline resulted in more SE events (**Fig 1B**, **Fig 3B**). However, morphine abstinence resulted in more SE events compared to methamphetamine abstinence (**Fig 1B**, **Fig 3B**).

DEU analysis on exonic regions across each phase of methamphetamine IVSA resulted in distinct DEU (515 exonic regions in MN vs SAL, 694 exonic regions in ABS vs SAL and 169 exonic regions in ABS vs MN) (FDR < 0.05 and |L2FC| > 0.5) and identified 1 exonic region that shared differential usage in all comparisons (FDR < 0.05 and |L2FC| > 0.5) (**Fig 3C**). Classification of exon usage throughout multiple phases of methamphetamine-taking and abstinence identified that reversible changes, or those that occur during methamphetamine-taking and return to baseline, was the most prevalent category (10 exonic regions in 9 genes) (**Fig 3D**). Both persistent and abstinence-specific exon dysregulation was identified in striatal microglia having undergone methamphetamine IVSA (10 exonic regions in 8 genes and 4 exonic regions in 4 genes, respectively) and morphine IVSA, however at a markedly lower level than in morphine IVSA (**Fig 3D**, **Fig 1E**).

### Methamphetamine and morphine induce differential splicing events in striatal microglia

To further evaluate the relationship between morphine and methamphetamine on AS in striatal microglia, we next compared the functional consequences of AS events occurring in both datasets. Predicted consequences were determined with transcript-aware functional annotation of rMATS splicing events. We focused on the 933 high confidence events occurring in genes that showed significant DEU (FDR < 0.05, |L2FC| > 0.5) and that contained significant AS events (FDR < 0.05, |ΔPSI| > 0.1) in each comparison. For each AS event, we determined the number of affected coding nucleotides by intersecting event coordinates with annotated coding sequences from gene-specific transcript models. Events were classified as frameshift-inducing when the affected coding nucleotides were not divisible by 3, predicting disruption of the downstream reading frame. Events with coding nucleotides divisible by 3 were classified as in-frame deletions (for SE’s) or in-frame changes (for MXE’s). Events without coding sequence overlap were classified as affecting noncoding or untranslated regions. 147 events that could not be reliably mapped were excluded, and 786 events were classified (622 from the morphine dataset and 164 from the methamphetamine dataset). Of these, 216 (27.5%) were predicted to cause frameshifts, 276 (35.1%) predicted to cause in-frame deletions, 96 (12.2%) predicted to result in in-frame changes, and 198 (25.2%) predicted to affect noncoding regions (**Fig 4A**). The general proportion of predicted functional consequences was similar across both data sets with in-frame deletions being the most common (34.1% in methamphetamine and 35.4% in morphine) (**Fig 4B**). While predicted frameshifts were distributed across all event types, SEs contributed the most predicted frameshift events (114), followed by RI’s (43), MXE’s (38), A3SS (15), and A5SS (6) (**Fig 4C**).

**Figure 4.**
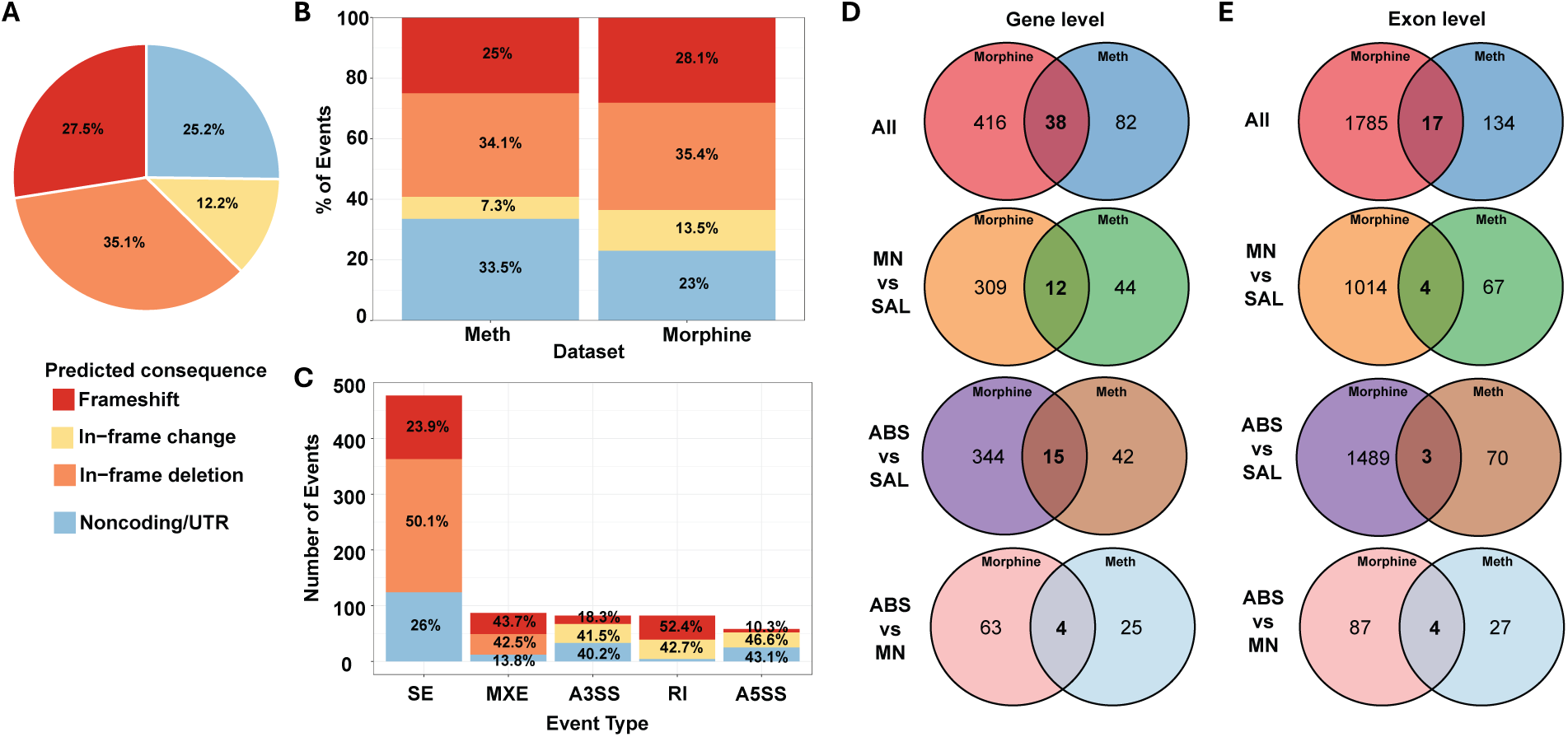
Predicted AS functional consequences are similar in OUD and MUD. **(A)** Distribution of predicted functional consequences for AS events classified by rMATS in both morphine and methamphetamine datasets. **(B)** Stacked bar plots showing the proportion of each predicted consequence in methamphetamine and morphine datasets. **(C)** rMATS AS event classification representing the contribution of each splicing event-type to an associated functional consequence. **(D)** Number of overlapping genes containing DEU event and AS events across pairwise comparisons in striatal microglia from morphine and methamphetamine datasets. **(E)** Number of overlapping DEU’s that also have AS events across pairwise comparisons in striatal microglia from morphine and methamphetamine datasets, regardless of L2FC direction.

Given that the main difference between the two datasets was the type of drug consumed, and that the predicted overall functional consequences were similar (**Fig 4A-C**), we next sought to identify the exonic regions that shared differential usage. To this end we overlapped high confidence events identified in striatal microglia for each pairwise comparison for both substances and considered these to be overlapping if the exonic region was significantly differentially used in the same comparison (FDR < 0.05, |L2FC| > 0.5). Interestingly, we did not find many overlapping DEUs between the datasets, with all comparisons yielding only 38 genes that contained any DEUs in both datasets and 17 exonic regions being differentially used (**Fig 4 D-E**). ABS vs SAL contained the most overlap with 15 genes containing at least one AS event and DEU in both data sets (**Fig 4D**) and MN vs SAL and ABS vs MN the most overlap with 4 DEUs shared between both datasets (**Fig 4E**). These data suggest that AS occurs in striatal microglia regardless of the drug consumed, whereas the genes impacted by AS machinery are indeed drug dependent.

## Discussion

Microglia are the primary immune cells of the CNS and play a key role in development and homeostasis, as well as in disease states^47^. In fact, microglial activity may depend on communication with neurons and other glia, which can induce transcriptional changes in microglia via several mechanisms, including transcriptional regulatory elements, chromatin accessibility and epigenetic modifications^48–50^. While alternative splicing events have been well characterized in colorectal, breast, stomach and neurological cancers^51–54^ and even in some SUDs^55–58^, studies on drug-induced AS events in microglia have remained limited. This is due in part to the fact that studies on microglia in SUDs have been limited to the transcriptional level. While these studies have yielded valuable insights into transcriptional responses of microglia, the underlying mechanisms by which transcripts switch isoforms and how AS impacts protein function remain unclear. Here we employed a clinically relevant model of drug-taking and utilized publicly available data to conduct DEU and AS event analysis on transcripts from striatal microglia. In doing so, we uncovered dysregulation of AS mechanisms in microglia during morphine-taking and abstinence as well as methamphetamine-taking and abstinence.

Specifically, we found that SE events are the most predominant AS events that occur across all tested conditions of both morphine and methamphetamine IVSA (**Fig 1B**, **Fig 3B**), which align with previous studies indicating that SE events are the most prevalent form of AS events in post-mortem human brains from patients with OUD^56^ as well as in the nucleus accumbens from methamphetamine treated mice^59^. Furthermore, we were able to identify and classify DEUs based on usage patterns, indicating that morphine induces persistent changes in exon usage (**Fig 1E**), while methamphetamine DEUs fit into more reversible patterns (**Fig 3E**). This is most likely a consequence of pharmacological differences between the two classes of drugs^60–64^. Interestingly, while we could not match many patterns between the two drugs, the predicted functional consequences of AS events were relatively similar with in-frame deletions being the most common predicted outcome across both data sets (**Fig 4A-C**). Secondary to in-frame deletions, for which functional protein consequences would be hard to predict, a large contribution of predicted frameshifts as a result of of AS events was also shared between the two datasets (**Fig 4B**). Further validation could reveal the implications of such a large number of predicted functional impacts. GO analysis of genes that featured persistent patterns of DEUs concomitant to AS events identified enrichment in RNA processing, chromatin organization, and protein stability pathways. Functional annotation of AS events within these pathways revealed that 27 genes harbor predicted frameshift-inducing events. Of these, several have related microglial functions, including autophagy (*Wdr81*), RNA processing (*Hnrnpl*, *Mbnl2*, *Tia1*), chromatin remodeling (*Chd4*), and cytoskeletal dynamics (*Abl1*, *Cd2ap*). *Wdr81*, which contained an A5SS event affecting 88 coding nucleotides, is involved in protein stability pathways (**Fig 2A-B**). This predicted frameshift in *Wdr81* could reduce functional protein levels via nonsense mediated decay (NMD), which degrades mRNA containing premature stop codons^65^. Impaired *Wdr81* function has been shown to disrupt autophagosome-lysosome fusion and when lost has been implicated in numerous neurodevelopmental disorders, including learning disabilities, cerebellar ataxia, and microcephaly^43^. In microglia, this loss of function could lead to accumulation of damaged organelles and debris in lysosomes leading to increased inflammatory signaling^66^. AS events in AS machinery genes such as *Hnrnpl*, a key regulator of AS which has been extensively studied in neurons^67,68^, are of high interest. *Hnrnpl* has been shown to be a key modulator of B-cell activation via regulation of genes such as lysine demethylase 6b (*Kdm6b)*^41^, an enzyme responsible for the erasure of the transcriptionally-repressive histone mark, H3K27me3. *Hnrnpl* has also been shown to regulate *Tnfα* gene expression via lncRNA mechanisms ^69^. The RI in *Hnrnpl* may represent a splicing factor that underwent aberrant splicing in response to morphine exposure, indicating a potential autoregulation of splicing in microglia. A frameshift could reduce functional Hnrnpl protein, potentially creating a feed-forward dysregulation of splicing programs and contributing to morphine-induced alterations in microglial activation states. Together, these findings suggest that morphine exposure may compromise multiple critical microglial proteins. Furthermore, the persistence of DEUs during morphine abstinence suggests establishment of long-lasting microglial dysfunction that may contribute to chronic neuroinflammatory states in OUD^70^.

While there were fewer pattern specific genes in methamphetamine than in morphine, the overall levels of shared AS events demonstrate that these events are altering microglial transcripts in both OUD and MUD. While we attempted to conduct functional classification, an important limitation is that we were unable to account for reading frame phase as well as other mechanisms such as NMD escape which could change the predicted outcomes. Our goal was not to predict the direct outcome of AS events, but instead to generate these broad classifications to enable future studies to build upon our findings and generate more robust parameters to better define functional outcomes. Future studies may also include analyses that consider gene expression to detect isoform switching which would complement these findings. Additionally, since the transcriptional consequences of drug-taking are known be sexually dimorphic,^71^ future studies should incorporate males and females to better understand such sex-specific effects. Taken together, the findings presented here highlight AS dysregulation as a potentially novel mechanism underlying opioid-induced and methamphetamine-induced microglial adaptations and identify potential candidates for follow-up studies.

## Methods

### Animals

Male C57BL/6J mice (12–16-week-old, ∼25-30 g; Jackson Laboratories, Bar Harbor, ME; SN: 000664) were housed in the animal facilities at the University of Miami Miller School of Medicine. Mice were maintained on a 12:12 h light/dark cycle and were housed three to five per cage. Animals were provided with food and water *ad-libitum*. All animals were maintained according to National Institutes of Health (NIH) guidelines, and animal experiments were performed in Association for Assessment and Accreditation of Laboratory Animal Care (AAALAC) accredited facilities. All experimental protocols were approved by the Institutional Animal Care and Use Committee (IACUC) at the University of Miami Miller School of Medicine.

### Jugular catheter surgery

Mice were prepared with indwelling jugular catheters as previously described^46,72,73^. Briefly, mice were anesthetized with an isoflurane (1–3%)/oxygen vapor mixture and prepared with indwelling jugular catheters. Briefly, the catheters consisted of a 6.5-cm length of Silastic tubing fitted to guide cannula (PlasticsOne, Protech International Inc., Boerne, TX, USA) bent at a curved right angle and encased in dental acrylic and silicone. The catheter tubing was passed subcutaneously from the animal’s back to the right jugular 14 vein, and 1-cm length of the catheter tip was inserted into the vein and anchored with surgical silk sutures. Mice were administered Meloxicam (5 mg/kg) subcutaneously prior to start of surgery and 24 hours post-surgery. Catheters were flushed daily with physiological sterile saline solution (0.9% w/v) containing heparin (10–60 USP units/mL) beginning 48 hours after surgery. Animals were allowed 3-5 days to recover from surgery before commencing intravenous morphine self-administration. Catheter integrity was tested with the ultra-short-acting barbiturate anesthetic Brevital (methohexital sodium, Eli Lilly, Indianapolis, IN, USA).

### Operant self-administration training

Morphine sulfate (NIDA Drug Supply Program, Research Triangle Park, NC, USA), used for self-administration, was dissolved in 0.9% sterile saline. Mice were trained and underwent self-administration as previously described ^46,73^. Briefly, mice were trained to self-administer morphine or methamphetamine during 2-hour daily sessions for 5 days at fixed ratio 3 (FR3) schedule reinforcement. Morphine (0.1 mg/kg/infusion; 36 μl/infusion; 3 second infusion) or methamphetamine (0.05 mg/kg/infusion; 14 μl/infusion; 2 second infusion)^46^ was delivered through Tygon catheter tubing (Braintree Scientific, MA, USA) into the intravenous catheter by a variable speed syringe pump (Med Associates Inc, Fairfax, VT, USA). Each self-administration session was performed using two retractable levers (one active, one inactive) that extended 1 cm into the chamber. Successful completion of criteria on the active lever was accompanied by a 20 second cue-light and time out (TO) period. Responses on the inactive lever were recorded but had no scheduled consequences, whereas the active lever was used to train mice from a fixed ratio of 1 to 3. After the initial 5 days (acquisition), mice were allowed to self-administer morphine (0.3 mg/kg/infusion; 36 μl/infusion; 3 second infusion) during 10 consecutive daily 2-hour maintenance sessions (FR3TO20). Animals that did not demonstrate stable responding (greater than 7 infusions, less than 25% variability, and a 2:1 ratio of active and inactive lever pressing across three sessions) or showed signs of compromised catheter patency were excluded from analysis. For forced home-cage abstinence, mice remained in their home cage for 21 days with no access to the drug.

### Microglial isolation

Microglia were isolated as previously described^46^, with minor modifications. Briefly, mice were anesthetized with isoflurane and perfused through the ascending aorta with 1X phosphate buffer saline (PBS; pH 7.4; ThermoFisher, 10010023) plus heparin (7,500 USP units). The striatum from individual mice were dissected at the indicated time points of the saline (SAL) (n = 3), maintenance (MN) (n = 4), and abstinence groups (ABS) (n = 4). Resulting tissue was then transported on ice in Hibernate A Medium (Gibco, A1247501) until dissociation on the gentleMACS Octo Dissociator (Miltenyi Biotec, #130-096-427) using the Adult Brain Dissociation Kit (Miltenyi Biotec, #130-107-677) according to manufacturer’s instructions. All steps after initial dissociation were performed on ice and all tubes were prechilled. The resulting single cell suspension was incubated with anti-mouse CD11b (microglia-specific) magnetic MicroBeads (Miltenyi Biotec, #130–093–634) and microglia were positively selected via column purification (Miltenyi Biotec, #130-042-201).

### RNA extraction and Next Generation Sequencing library preparation

The eluted fraction containing purified microglia was then added to 350 µL RLT plus buffer (Qiagen, 1053393) for extraction and purification of total RNA according to manufacturer’s instructions using the Qiagen AllPrep DNA/RNA Mini Kit (Qiagen, 80204). Total RNA input was normalized and NGS libraries were prepared using NEBNext Single Cell/Low Input RNA Library Prep Kit for Illumina (New England BioLabs, E6420S) according to the manufacturer’s instructions. Sequencing was performed on an Illumina NovaSeq6000 platform (150×150bp PE) targeting 30 million reads per sample by Azenta Life Sciences.

### RNA-sequencing differential gene expression pre-processing

Prior to mapping, raw RNA-seq datasets were first trimmed using TrimGalore (v.0.6.7)^74^with cutadapt (v.1.18)^75^. Illumina sequence adaptors were removed, the leading and tailing low-quality base-pairs (fewer than 3) were trimmed. Next, reads with a length of at least 20-bp were mapped to the genome using STAR (v.2.7.10a)^76^ with the following parameters: –outSAMtype BAM SortedByCoordinate –outSAMunmapped Within –outFilterType BySJout –outSAMattributes NH HI AS NM MD XS –outFilterMultimapNmax 20 –outFilterMismatchNoverLmax 0.3 --quantMode TranscriptomeSAM GeneCounts. The resulting bam files were then passed to StringTie (v.2.1.7)^77^ to assemble sequenced alignments into estimated transcript and gene count abundance given the Gencode GRCm38 (NCBI) transcriptome assembly.

### Differential gene expression analyses

The R/Bioconductor DESeq2 (v.1.34.0)^78^ package was used to detect the differentially expressed genes between striatal microglia. Only genes meeting independent filtering criteria as determined by DESeq2, and adjusted *p*-value (padj) < 0.05 were considered as differentially expressed and were used for downstream analysis.

### Alternative splicing detection analysis

Alternative splicing (AS) events were identified using rMATS (v4.1.2)^38^ with junction count-based analysis. rMATS was run separately for two experimental paradigms: morphine IVSA and methamphetamine IVSA. For each drug of interest, morphine and methamphetamine, three pairwise comparisons were performed to assess splicing changes: MN vs SAL, ABS vs SAL and ABS vs MN. For each comparison, rMATS detected and quantified five types of AS events: skipped exons (SE), mutually exclusive exons (MXE), alternative 3’ splice sites (A3SS), alternative 5’ splice sites (A5SS), and retained introns (RI). Significance was determined using a false discovery rate (FDR) < 0.05 for initial plots. Downstream analysis included filtering based on absolute inclusion level difference using changes in percent spliced in (PSI) (|ΔPSI| > 0.1). PSI values were calculated from junction-spanning read counts to quantify the relative usage of alternative splice forms. Significant splicing events were annotated with gene symbols and event types.

### Differential exon usage analysis

Differential exon usage (DEU) was analyzed using DEXSeq (v1.42.0)^39^ to identify exonic regions with condition-specific changes in relative usage independent of overall gene expression. Analysis was performed separately for morphine and methamphetamine studies. DEXSeq exon counting bins were generated from NCBI RefSeq annotations (mm10/GRCm38) using dexseq_prepare_annotation.py^39^. Two chromosome Y genes (Erdr1, G530011O06Rik) with problematic overlapping features were excluded from the annotation prior to flattening to prevent coordinate conflicts during bin generation. Pairwise DEXSeq analyses were conducted as they were done with the rMATS analyses described above. Each comparison was analyzed independently with comparison-specific dispersion estimates and statistical tests to obtain comparison specific L2FC’s and padj values. Size factor estimation was done using the median-of-ratios method, dispersion estimation was conducted using a generalized linear model with exon-by-condition interaction terms and a likelihood ratio test was used to test for differential exon usage. Fold change estimations were generated for each exonic region relative to other exons in the same gene and multiple testing correction was performed using the Benjamini-Hochberg procedure. Exons were considered significantly differentially used if they met FDR < 0.05 and |L2FC| > 0.5.

### Identification of high confidence AS events

To identify high confidence AS events, DEXSeq and rMATS analyses were integrated at the gene level. DEXSeq exons showing significant differential usage (FDR < 0.05, |L2FC| > 0.5) were retained only if their parent gene also harbored at least one significant rMATS AS event in the same comparison (FDR < 0.05, |ΔPSI| > 0.1). This filtering was performed at the gene level rather than requiring coordinate-level correspondence between DEXSeq bins and rMATS events, providing validation that genes with DEU may exhibit independent evidence of AS. The resulting exon sets were used for pattern classification and cross-dataset comparisons.

### Exon usage pattern classification

Events were classified by comparison to identify temporal patterns of splicing regulation during drug-taking and abstinence. Significant exons were classified into temporal response patterns based on their significance profiles across the three comparisons: Reversible: drug-induced changes (significant exon usage change in MN vs SAL comparisons) that normalize after drug-abstinence (non-significant change in exon usage in ABS vs SAL, significant exon usage change in ABS vs MN). Persistent: drug-induced changes (significant exon usage change and exon usage L2FC in the same direction in MN vs SAL and ABS vs SAL) that remain after drug-abstinence (non-significant change in exon usage in ABS vs MN). Abstinence-Specific: Changes emerging only after forced home cage abstinence (non-significant change in exon usage in MN vs SAL, significant exon usage change in ABS vs SAL and ABS vs MN significant). Progressive: effects that continue or amplify post-withdrawal (all three comparisons show significant exon usage change, and the ABS change is larger than MN). Rebound: paradoxical overcorrection after drug abstinence (all three comparisons showed significant exon usage change, but L2FCs were in opposite directions). Partial-Reversal: incomplete normalization toward baseline. Exons that exhibited significance in at least one comparison but did not fit a clear pattern described above were not classified.

### Functional prediction classification of exons

Function annotation was conducted on rMATS AS events occurring in genes with evidence of differential exon usage. Specifically, we included rMATS events (FDR < 0.05, |ΔPSI| > 0.1) from genes that also showed at least one exon with significant differential usage by DEXSeq (FDR < 0.05, |L2FC| > 0.5) in the same comparison. For each AS event, we identified the set of annotated transcripts for the parent gene and extracted coding sequence (CDS) coordinates from the Ensembl gene annotation (mm10). Transcripts were filtered to include only gene-specific transcript sets. Functional consequences were predicted using event-type-specific parameters. Events were classified as causing frameshifts or in-frame changes based on the number of affected coding nucleotides. For SEs, if the number of nucleotides overlapping the SE were not divisible by 3, they were classified as frameshift, otherwise they were classified as in-frame deletions. For MXEs the difference in length between alternative exons determined the outcome following the same description as SEs. For RIs and both ASS event types, the same divisibility rule was applied to the number of coding nucleotides in the affected region. Events were assigned to “Frameshift” if the affected coding nucleotides were not divisible by 3, predicted to shift the reading frame and likely introduce premature termination codons, “In-frame deletion” if SEs or shortened isoforms with coding nucleotides were divisible by 3, implying that the reading frame remained intact, “In-frame change” if ME exons had a length difference divisible by 3, predicted to be substituting one amino acid sequence for another, and “Noncoding/UTR” for events with no overlap with coding sequences, affecting only untranslated regions or non-coding transcripts. Events with genomic coordinates that could not be properly mapped to the reference genome or transcript annotations were excluded. Therefore, all proportional analyses, such as the percentage of frameshift vs. in-frame events, were calculated only from events with valid functional annotations.

### Functional enrichment analysis

For DEG analyses, the R/Bioconductor clusterProfiler (v.4.2.2)^79,80^ package was used to perform GO analysis on differentially expressed genes using a background of all genes with detectable expression. Only the GO terms and pathways with a pvalue < 0.05 and qvalue < 0.1 following false discovery rate (FDR) correction were considered, with focus given to those containing genes and pathways related to AS. Specifically GO terms were selected if they contained the following terms: “splic”, “spliceosome”, “snrnp”, “intron”, “exon”, “exon junction”, “mRNA processing”, “pre-mRNA|premrna”, “alternative splicing” or “RNA splicing”. The associated GO and pathway enrichment plots were generated using the ggplot2 package (v.3.4.2)^81^.

For DEU analyses, the R/Bioconductor clusterProfiler (v.4.2.2)^79,80^ package was used to perform GO analysis on high confidence splicing genes in the persistent category, using a background of all genes with detected exonic expression. Only the GO terms and pathways with a pvalue < 0.05 and qvalue < 0.05 following a Benjamini-Hochberg (BH) procedure to control for FDR were considered. The associated GO and pathway enrichment plots were generated using the ggplot2 package (v.3.4.2)^81^. All the other plots were generated using the ggplot2 package^81^.

### GEO data sets and processing

FATSQ files were downloaded from the Gene Expression Omnibus (GEO) data repository from GSE245951. The files were then processed using the same methods as described above.

### Statistical analysis

Statistical analyses were conducted in accordance with the various packages used for RNA-sequencing and AS data. Specifically, stringent cutoffs were used for significance detection in differential gene expression analysis (padj < 0.05), differential exon usage (FDR < 0.05, |L2FC| > 0.5) and AS detection (FDR < 0.05, |ΔPSI| > 0.1). Information on specific analyses run can be found in the associated code and methods sections.

### Support

This work was supported by NIH grants DP1DA051828 (LMT), R01AA029924 (LMT), F30DA060666 (LLB), and K00DA062570 (SJV) as well as by a kind gift from the Shipley Foundation (LMT)

## Author contributions

Conceptualization: AVM, LMT Methodology: AVM, SJV, LLB, LMT Software: AVM

Investigation: AVM, SJV, LLB Visualization: AVM

Data Curation: AVM, LMT Supervision: LMT Writing—original draft: AVM, LMT

Writing—review & editing: AVM, SJV, LMT

## Data and materials availability

All data files and code used in processing and analysis will be available from GEO/SRA and GitHub repositories upon publication and/or upon request.

